# Functional diversity of phage sponge proteins that sequester host immune signals

**DOI:** 10.1101/2025.08.24.671296

**Authors:** Romi Hadary, Renee B. Chang, Nathalie Béchon, Nitzan Tal, Ilya Osterman, Erez Yirmiya, Jeremy Garb, Gil Amitai, Philip J. Kranzusch, Rotem Sorek

## Abstract

Multiple bacterial immune systems, including CBASS, Thoeris, and Pycsar, employ signaling molecules that activate the immune response following phage infection. Phages counteract bacterial immune signaling using sponge proteins that bind and sequester the immune signals, but the breadth of immune signals targeted by phage sponges is unclear. Here we study the functional versatility of Acb2, Tad1 and Tad2, three families of sponge proteins known to inhibit CBASS and Thoeris signaling. Eighty-four proteins representing the phylogenetic diversity of these sponge families were tested for their ability to inhibit immunity by sequestering 3′3′-cGAMP and 3′3′-cUA (CBASS), cCMP and cUMP (Pycsar), and 3′cADPR, His-ADPR and N7-cADPR (types I, II and IV Thoeris, respectively). While Acb2 proteins were so far reported to inhibit only CBASS systems, we found Acb2 homologs that bind 3′cADPR and inhibit Thoeris defense. In addition, we discovered sponge proteins that inhibit Pycsar and type IV Thoeris by binding cUMP and N7-cADPR, respectively. Using crystal structures, structural modeling and biochemical analyses, we explain the molecular basis for signal-binding specificities in members of these sponge families. Our study reports the first sponges inhibiting Pycsar and type IV Thoeris, and demonstrates how phage sponges evolve to obtain diverse specificities.

## Introduction

Defense systems that rely on small molecule signaling are emerging as important constituents of the bacterial anti-phage defense arsenal^1–11^. These systems typically comprise two main components, one that recognizes phage infection and generates the immune signaling molecule, and another that senses the signaling molecule and executes the immune response^12^.

Bacterial immune signaling systems include CBASS^1^, Pycsar^3^ and Thoeris^2^. The hallmark of CBASS systems are enzymes of the CD-NTase family, which produce cyclic di-nucleotide or cyclic tri-nucleotide signals in response to infection^1,9,13^. CBASS is the evolutionary ancestor of the human cGAS-STING antiviral pathway, which uses 2′–5′ / 3′–5′ cyclic GMP-AMP (cGAMP) as an immune signaling molecule^1,14–16^. The Pycsar system encodes pyrimidine cyclase enzymes that generate either 3′,5′ cyclic cytidine monophosphate (cCMP) or 3′,5′ cyclic uridine monophosphate (cUMP) as immune signaling molecules^3^. The signal producing enzymes in Thoeris systems comprise TIR domains that process the molecule NAD^+^ to form signaling molecules that include adenine diphosphate ribose (ADPR) moieties^2,4,5^. Thoeris is probably ancestral to a central immune pathway in plants in which TIR domain proteins produce ADPR-containing signaling molecules that activate downstream immunity^1,17^.

It was estimated that at least 20% of all bacteria encode immune signaling systems^18^. Phages, on their end, have evolved molecular mechanisms to evade or suppress these immune systems. A central phage evasion strategy involves sequestration of the immune signaling molecule using dedicated phage proteins called “sponges”. These are typically small proteins that form homo-oligomeric complexes, each containing several pockets capable of sequestering multiple immune molecules with high affinity^19–21^. The first discovered sponge was Thoeris anti defense 1 (Tad1), a protein from the *Bacillus* phage SBSphiJ7 that inhibits type I Thoeris by binding and sequestering 3′cADPR (also called 1′′–3′ gcADPR), the immune signaling molecule produced by this system^19^. Tad2 is another protein, unrelated in sequence or structure to Tad1, which also inhibits type I Thoeris by binding 3′cADPR^19,20^. A third family of sponge proteins, called Anti-CBASS 2 (Acb2), bind and sequester cyclic di-nucleotides and cyclic tri-nucleotides to inhibit CBASS signaling^21,22^. Recent findings suggest that additional families of phage-encoded sponge proteins remain to be discovered^23,24^.

Recent studies show that homologs within the same sponge family can differ in their binding specificities to immune signaling molecules. Specifically, while Tad2 from phage SPO1 binds 3′cADPR to inhibit type I Thoeris, a homolog of Tad2 from *Myroides odoratus* prophage (*Mo*Tad2) sequesters the signaling molecule of type II Thoeris, comprising histidine conjugated to ADPR (His-ADPR)^5,20^. Additionally, a Tad2 homolog derived from human gut metagenomic libraries (HgmTad2) can bind cyclic di-nucleotides in addition to binding 3′cADPR, and homologs of Tad1 can sequester, in addition to 3′cADPR, also CBASS-generated molecules such as 3′3′ cGAMP and the cyclic tri-nucleotides cAAA and cAAG^25^. These studies, based on a limited set of sponge homologs, show that phage sponge proteins can evolve to adapt their specificities and target diverse immune signaling molecules, but the breadth of this phenomenon, and the full complexity of the sponge roles in countering bacterial defenses, is currently unclear.

In this study we set out to explore the evolutionary and functional diversity of phage sponge proteins in nature. For this, we reconstructed the phylogenetic trees of the Tad1, Tad2 and Acb2 families using nearly 8,600 sponge protein homologs from phage and prophage genomes. We then experimented with 84 proteins, selected to span the sequence diversity of these sponges, and tested their capacity to inhibit seven variants of CBASS, Pycsar and Thoeris systems that use diverse signaling molecules. This led to the discovery of the first sponges to inhibit Pycsar and type IV Thoeris, as well as to the revelation that Acb2, originally discovered as an anti-CBASS protein family, has a broader function and also inhibits Thoeris defense. Our study documents remarkable functional plasticity for phage sponge proteins, and identifies the structural and molecular basis for diversification of their signal binding properties.

## Results

To study the functional diversity of counter-defense sponge families, we first set out to construct phylogenetic trees for each family. For this, we queried the IMG/VR v3 database of phage proteins for homologs of Acb2, Tad1 and Tad2, yielding 4,204 non redundant homologs of Acb2, 1,587 of Tad1, and 2,783 Tad2 homologs (Methods) (Table S1)^26,27^. We then reconstructed phylogenetic trees for each of the three families, and selected homologs spanning the diversity of each family for experimental examination. Overall, 84 sponge proteins were cloned for further experiments, including 46 Acb2, 20 Tad1, and 18 Tad2 proteins (Table S2).

Each of the selected sponge proteins was introduced into cells expressing one of seven defense systems that rely on immune signaling molecules. These included CBASS from *Escherichia albertii* MOD1-EC1698 (producing 3′3′-cGAMP)^28^, CBASS from *Yersinia aleksiciae* IP28587 (3′3′-cUA)^28^, Pycsar from *Escherichia coli* E831 (cCMP)^3^, Pycsar from *Bacillus* sp. G1 (cUMP)^24^, type I Thoeris from *Bacillus cereus* MSX-D12 (3′cADPR)^2,19^, type II Thoeris from *Bacillus amyloliquefaciens* Y2 (His-ADPR)^5^, and type IV Thoeris from *Pseudomonas* sp. 1-7 (N7-cADPR)^4^. These systems were integrated into the genomes of *B. subtilis* BEST7003 or *E. coli* MG1655 under the control of their native promoters (for *B. subtilis*) or under a promoter inducible by anhydrotetracycline (aTc) (for *E. coli*).

We used a previously reported high-throughput screening methodology to test the effect of the 84 selected sponge proteins on these defense systems^24^. In this method, sponge gene transformations are performed in liquid cultures in a 96-well plate format, and these cultures are then subjected to phage infection at low multiplicity of infection (MOI) using a phage naturally restricted by the defense system. Cultures in which the defense system is active continue to grow, while cultures where the sponge protein inhibited defense collapse due to phage propagation, allowing to record sponge activity *in vivo* using optical density measurements. Cases in which sponge activity was observed in liquid cultures were further confirmed via plaque assays.

Overall, 65 of the 84 tested proteins exhibited activity against at least one defense system, with 25 proteins inhibiting more than one of the tested systems (Table S2). Most Acb2 proteins inhibited one or both of the CBASS systems in our set, as expected from prior studies on Acb2^21,22,29^. Surprisingly, however, a substantial fraction of Acb2 proteins inhibited type I Thoeris, suggesting that the Acb2 sponge family is not strictly limited to CBASS inhibition. Proteins from the Tad1 family were able to inhibit both CBASS and type I Thoeris defense, in agreement with previous reports^19,25^. The Tad2 protein family showed remarkable functional diversity, as CBASS, Pycsar and the three types of Thoeris systems tested were inhibited by at least one Tad2 homolog, with many Tad2 proteins inhibiting more than one system.

To understand the structural and biochemical determinants allowing sponges from the same family to inhibit different defense systems, we selected a subset of these sponges for deeper characterization. Specifically, we studied Acb2 sponges that inhibit type I Thoeris, Tad2 sponges that inhibit Pycsar, and Tad2 sponges inhibiting type IV Thoeris. These are described in the following sections.

### Acb2 homologs inhibit type I Thoeris

We identified 10 Acb2 homologs that inhibited type I Thoeris defense when co-expressed with the defense system in vivo (Fig. 1A-C). To test if these proteins are capable of inhibiting Thoeris in more native settings, i.e., when expressed from a phage genome, we engineered one of the Thoeris-inhibiting Acb2 homologs (Acb2_43) into the genome of SBSphiJ, a phage typically restricted by type I Thoeris^2^. SBSphiJ phage carrying *acb2_43* under the control of a native *tad1* promoter showed improved infectivity on Thoeris-expressing cells, suggesting that this protein natively inhibits Thoeris (Fig. 1D).

**Figure 1.**
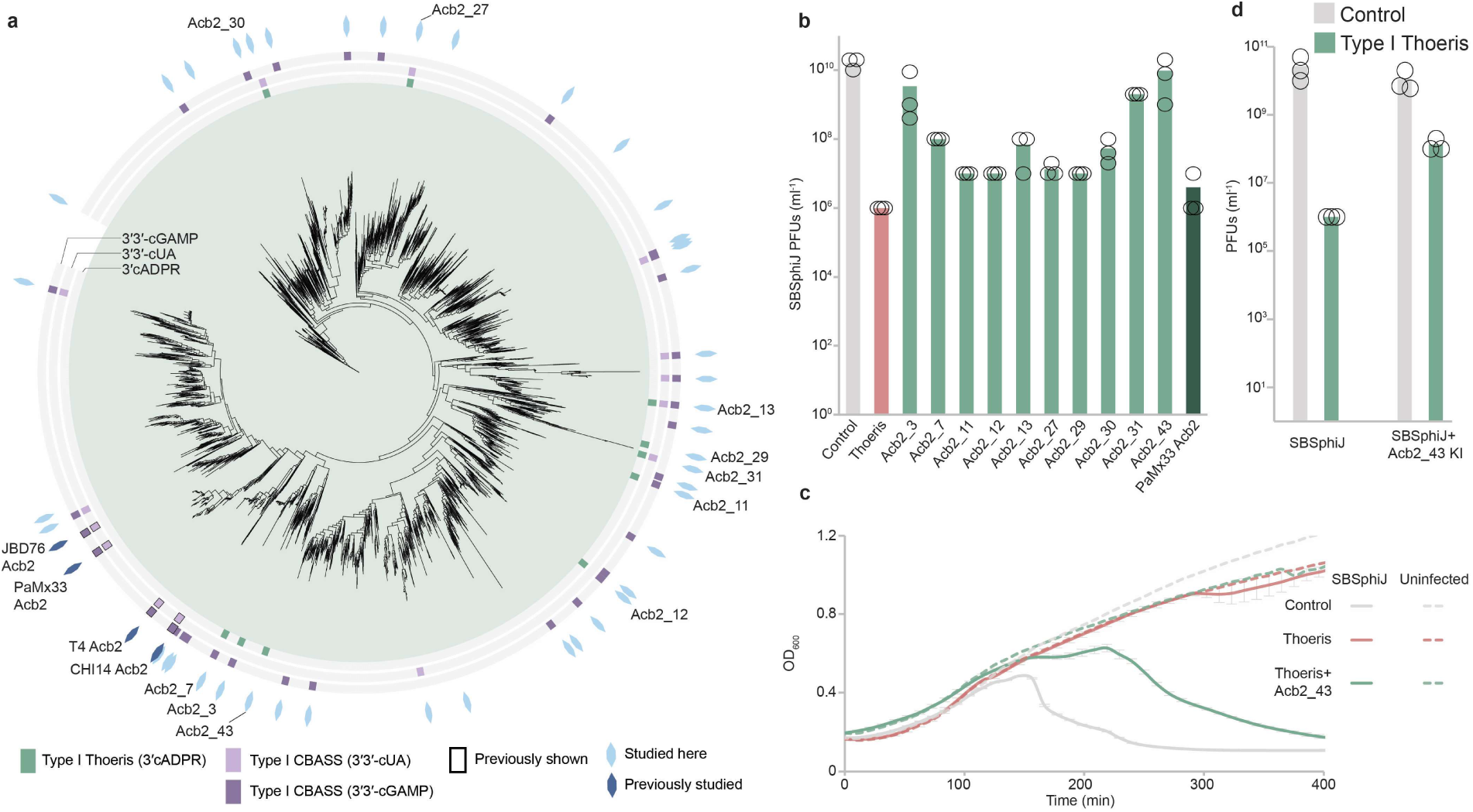
**Acb2 homologs inhibit Type I Thoeris**. (A) Phylogenetic analysis of Acb2 homologs from phage genomic and metagenomic protein databases (n = 4,204 non redundant sequences). Previously studied homologs are marked by dark blue hexagons on the outer rim of the tree; homologs tested in the current study are in light blue. Homologs that inhibit the defense system are marked with the respective color in the inner rings. (B) Plating efficiency of phage SBSphiJ on negative control *B. subtilis* cells (no system, grey), cells expressing type I Thoeris (red), and cells co-expressing type I Thoeris and a homolog of Acb2 (green). Data represent PFUs per milliliter, and bars show the average of three replicates with individual data points overlaid. (C) Growth curves of *B. subtilis* cells expressing type I Thoeris (red), co-expressing type I Thoeris and an Acb2 homolog (green) or control cells (grey). Cells were infected with SBSphiJ phage at an MOI 0.1. Each curve is the average of three replicates, with error bars indicating standard deviation, except for uninfected control (one replicate). (D) A phage engineered to express the Acb2 homolog Acb2_43 overcomes Thoeris defense. Shown are data for wild-type SBSphiJ phage, as well as SBSphiJ knocked-in (KI) with Acb2_43. Data represent PFUs per milliliter of phages infecting control cells (no system), or cells expressing the type I Thoeris system. Bars show the average of three biological replicates with individual data points overlaid.

Type I Thoeris comprises ThsB, a TIR-domain protein that produces a 3′cADPR signaling molecule in response to infection, and ThsA, an NADase whose activity is triggered by the signaling molecule^2,19^. To verify that inhibition of type I Thoeris by Acb2 was associated with elimination of the Thoeris signaling molecule in infected cells, we collected lysates from cells expressing ThsB. These lysates trigger the NADase activity of purified ThsA in vitro, because ThsB produces 3′cADPR in infected cells. However, lysates from ThsB-expressing cells that co-expressed Acb2 proteins failed to activate ThsA in vitro, suggesting that these Acb2 homologs sequestered the signaling molecule produced by ThsB (Fig. 2A).

**Figure 2.**
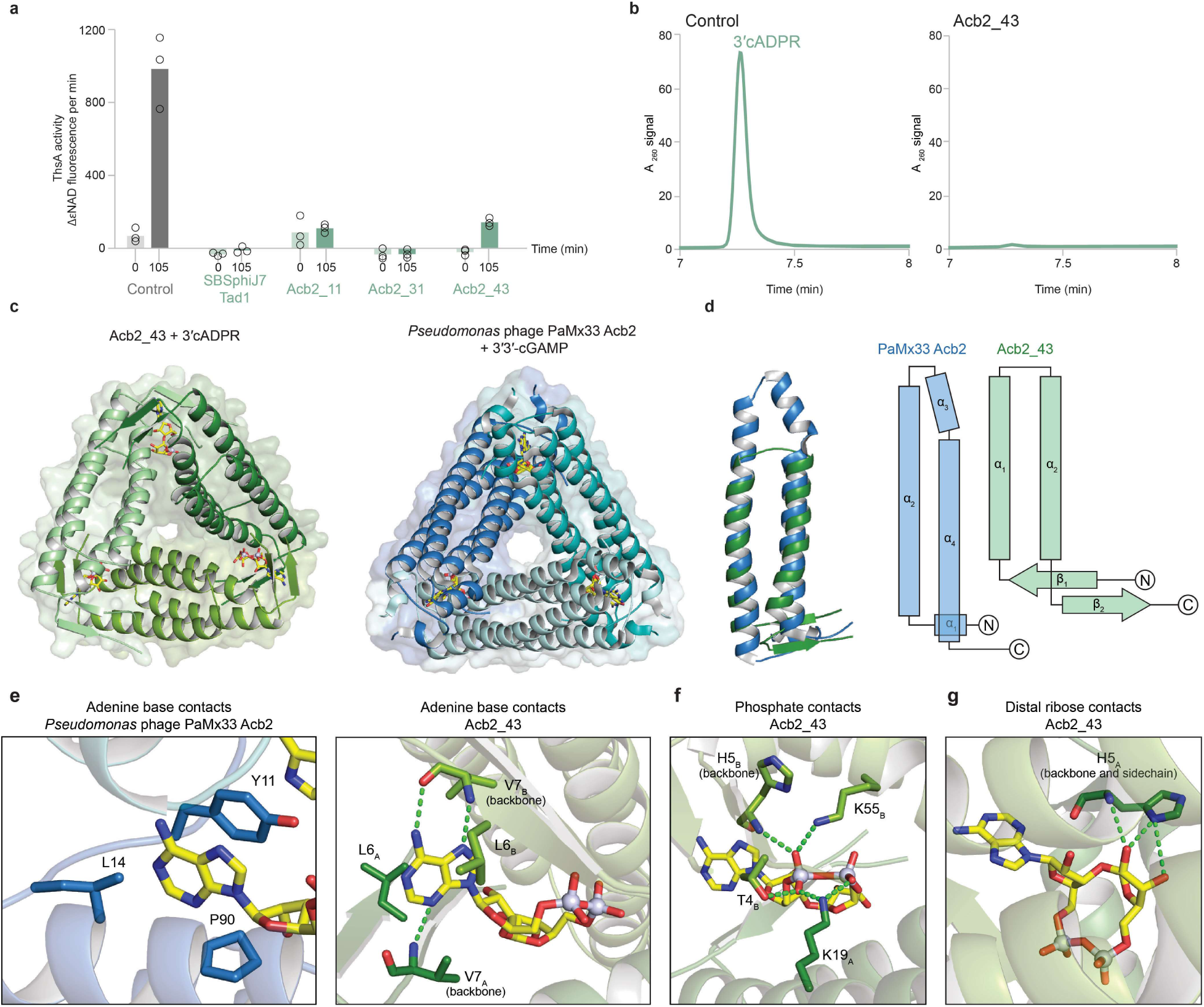
Molecular basis for suppression of Type I Thoeris by Acb2 homologs. (A) Acb2 proteins eliminate 3′cADPR in Thoeris-expressing infected cells. Cells expressing ThsB from *B. cereus* MSX-D12, or co-expressing both ThsB and an Acb2 homolog, were infected with phage SBSphiJ at a multiplicity of infection (MOI) of 10. Cell lysates were extracted before infection (t=0) and 105 minutes post-infection, and were filtered to retain small molecules. Filtered lysates were then incubated with ThsA, and the NADase activity of ThsA was measured using a nicotinamide 1,N6-ethenoadenine dinucleotide (εNAD) cleavage fluorescence assay. Bars represent the mean of three experiments, with individual data points overlaid. Control represents experiments with lysates from cells co-expressing ThsB and GFP. (B) HPLC analysis of 3′cADPR incubated with either purified GFP (control, left lane) or with purified Acb2_43 at 1:4 molar ratio. The data show that Acb2_43 eliminates 3′cADPR from the solution. (C) Overview of the crystal structure of Acb2_43 in complex with 3′cADPR, compared to previously determined structure of *Pseudomonas* phage PaMx33 Acb2 hexamer in complex with 3′3′-cGAMP (PDB: 8H2J). (D) Comparison and topology maps of *Pseudomonas* phage PaMx33 Acb2 (blue) and Acb2_43 (green) monomers. (E) Detailed view of PaMx33 Acb2 residues that interact with adenine base of 3′3′-cGAMP (left) and view of Acb2_43 residues that interact with adenine base of 3′cADPR (right). Green dashes indicate hydrogen bonding interactions and subscript denotes protomer chain. (F) Detailed view of Acb2_43 residues that interact with 3′cADPR diphosphate backbone. (G) Detailed view of Acb2_43 residues that interact with 3′cADPR distal ribose.

Incubation of a purified Acb2_43 with 3′cADPR in vitro resulted in elimination of the signaling molecule from the solution, showing that this protein efficiently sequesters the molecule (Fig. 2B). To define the molecular mechanism of Acb2_43 signal sequestration, we next determined a 1.6 Å X-ray crystal structure of Acb2_43 in complex with 3′cADPR (Fig. 2C). Acb2_43 adopts an overall architecture similar to previously characterized Acb2 family proteins and forms a hexameric assembly (Fig. 2C). Six protomers of Acb2_43 interlock to form a triangular, prism-like assembly, with distinct binding pockets located at each of the three vertices (Fig. 2C). Consistent with other members of the Acb2 sponge family, each Acb2_43 protomer contains a central region composed of two long anti-parallel helices, α_1_ and α_2_ (Fig. 2D). Interestingly, we observed that Acb2_43 protomers contain two additional β-strands at the N- and C-termini (β_1_ and β_2_). Residues from the β_1_-strand participate in forming the molecule-binding pocket of Acb2_43, suggesting that these structural elements may have evolved to enable Acb2_43 to sequester Thoeris signaling molecules (Fig. 2D).

In our experiments, Acb2_43 did not inhibit CBASS (Fig. S1A,B). Comparative analysis of the binding pockets of *Pseudomonas* phage PaMx33 Acb2, which inhibits CBASS defense and binds 3′3′-cGAMP, and Acb2_43, which inhibits Thoeris immunity and sequesters 3′cADPR, further underscores the diversity of Acb2 family proteins (Fig. 2E). In PaMx33 Acb2, the 3′3′-cGAMP adenine nucleobase is stabilized by π–π stacking interactions from residue Y11, and hydrophobic interactions from L14 and P90 (Fig. 2E). In contrast, Acb2_43 coordinates the adenine nucleobase of 3′cADPR through a distinct mechanism, where residues L6 and V7 on the β_1_ strand of two opposing protomers (Acb2_43_A_ and Acb2_43_B_, respectively) stabilize the base via a network of polar contacts and hydrophobic interactions (Fig. 2E). Similarly, the 3′cADPR diphosphate backbone is coordinated by contacts from the backbone of H5_B_ and sidechain-specific contacts from K19_A_, T4_B_, and K55_B_ (Fig. 2F). Finally, Acb2_43 residue H5_A_ anchors the ligand binding pocket and forms polar contacts with the 3′cADPR distal ribose (Fig. 2G). Together, these results illustrate the remarkable functional diversity of Acb2 sponge family proteins, and suggest that Acb2 proteins may have acquired molecular adaptions as a response to host signal diversification.

### Tad2 sponges that sequester Pycsar molecules

Two Tad2 homologs, Tad2_70 and *Mo*Tad2, were able to inhibit the Pycsar systems tested in our assays, suppressing both the Pycsar from *E. coli* E831 that produces cCMP, and the one from *Bacillus* sp. G1 producing cUMP (Fig. 3A,B, S2A). Interestingly, *Mo*Tad2 is a sponge that was previously shown to inhibit type II Thoeris by sequestering His-ADPR^5^. Engineering phage T5 to encode the sponge under the control of an early gene promoter resulted in a phage that can overcome the *E. coli* Pycsar, confirming the anti-Pycsar activity of the *Mo*Tad2 sponge (Fig. 3C).

**Figure 3.**
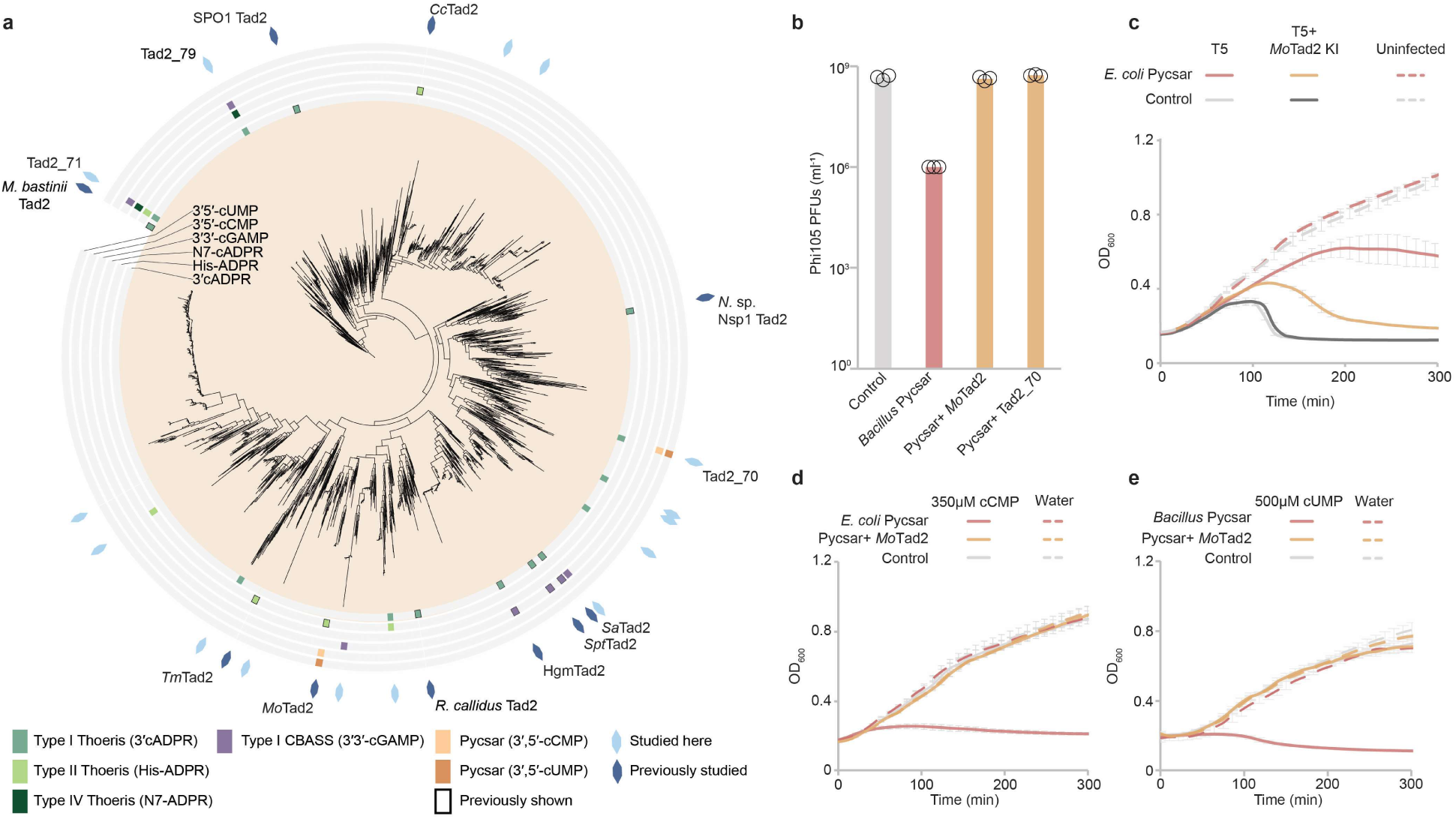
**Tad2 homologs inhibit Pycsar**. (A) Phylogenetic analysis of Tad2 homologs from phage genomic and metagenomic protein databases (n = 2,783 non redundant sequences). Previously studied homologs are marked by dark blue shapes on the outer rim of the tree; homologs tested in the current study are in light blue. Homologs that inhibit the defense system are marked with the respective color in the inners rings. (B) Plating efficiency of phage phi105 on negative control *B. subtilis* cells (no system, grey), cells expressing the Pycsar system from *Bacillus* sp. G1 (red), and cells co-expressing both Pycsar and a homolog of Tad2 (orange). Data represent PFUs per milliliter, and bars show the average of three replicates with individual data points overlaid. (C) Growth curves of *E. coli* cells expressing the Pycsar system from *E. coli* E831, or control cells not expressing Pycsar. Cells were infected with a wild-type T5 phage or with T5 phage engineered to express *Mo*Tad2 from an early phage promoter. Phages were infected at an MOI of 0.1. Each curve is the average of three replicates, with error bars indicating standard deviation. (D) Growth curves of *B. subtilis* cells expressing the Pycsar system from *Bacillus sp*. G1 (red), co-expressing Pycsar with *Mo*Tad2 (orange), or negative control cells (grey), with and without the addition of cUMP to the medium. Each curve represents the average of three replicates; error bars indicate standard deviation. (E) Growth curves of *E. coli* MG1655 cells expressing the Pycsar system from *E. coli* E831 (red), co-expressing Pycsar with *Mo*Tad2 (orange), or negative control cells (grey), with and without the addition of cCMP to the medium. Each curve represents the average of three replicates; error bars indicate standard deviation.

It was previously shown that growth of Pycsar-expressing cells is attenuated when 3′,5′ cCMP or 3′,5′ cUMP are added in high concentrations to the growth medium, most probably because sufficient amounts of the molecule penetrate the cell and activate the Pycsar effector^3^. In the current study we verified that growth of cells expressing the *E. coli* Pycsar (utilizing cCMP as the signaling molecule) was impaired when cCMP was added to the medium, and growth of *B. subtilis* cells expressing the *Bacillus* Pycsar was inhibited in the presence of cUMP (Fig. 3D,E). This growth inhibition effect was specific to the Pycsar signaling molecule, because the addition of non-cognate molecules, and specifically 3′,5′-cUMP to *E. coli* Pycsar or 2′,3′-cUMP to *Bacillus* Pycsar (a molecule different from the Pycsar signaling molecule 3′,5′-cUMP) did not affect growth of Pycsar-expressing cells (Fig. S2B,C). Notably, growth of Pycsar-expressing cells was not affected by supplementing cCMP/cUMP to the growth media when these cells also co-expressed *Mo*Tad2 (Fig. 3D,E). These results demonstrate that *Mo*Tad2 can rescue Pycsar cells from toxicity caused by cCMP or cUMP, further supporting that this sponge sequesters the Pycsar signaling molecules.

Recent structural studies showed that *Mo*Tad2 forms a homotetrameric structure and sequesters His-ADPR in two symmetric pockets, one located in the interface between protomers *Mo*Tad2_A_–*Mo*Tad2_B_, and the other in the *Mo*Tad2_C_–*Mo*Tad2_D_ interface^5^. However, introducing mutations in *Mo*Tad2 residues essential for the formation of this pocket resulted in a protein that was still able to inhibit the *Bacillus* Pycsar, implying that cUMP may be bound in a different location in that protein (Fig. 4A-C). Indeed, using AlphaFold3^30^ to co-fold a *Mo*Tad2 tetramer with cUMP resulted in a high-confidence structure model in which the molecule was predicted to be bound in a different site of the tetramer, formed at the interface between protomers *Mo*Tad2_A_–*Mo*Tad2_C_ (Fig. 4D). Point mutations in residues predicted by the AlphaFold3 model to be in the vicinity of cUMP in that pocket rendered *Mo*Tad2 unable to inhibit the *Bacillus* Pycsar (Fig. 4C,E) but retained its activity against type II Thoeris^5^. These results demonstrate that *Mo*Tad2 binds cUMP and His-ADPR in two separate pockets, explaining its ability to inhibit both Pycsar and type II Thoeris. Notably, the Tad2_A_–Tad2_C_ interface, which in *Mo*Tad2 forms the pocket that binds cUMP, was previously shown in another Tad2 homolog (HgmTad2) to bind 3′3′-cGAMP and other CBASS-related molecules^25^.

**Figure 4.**
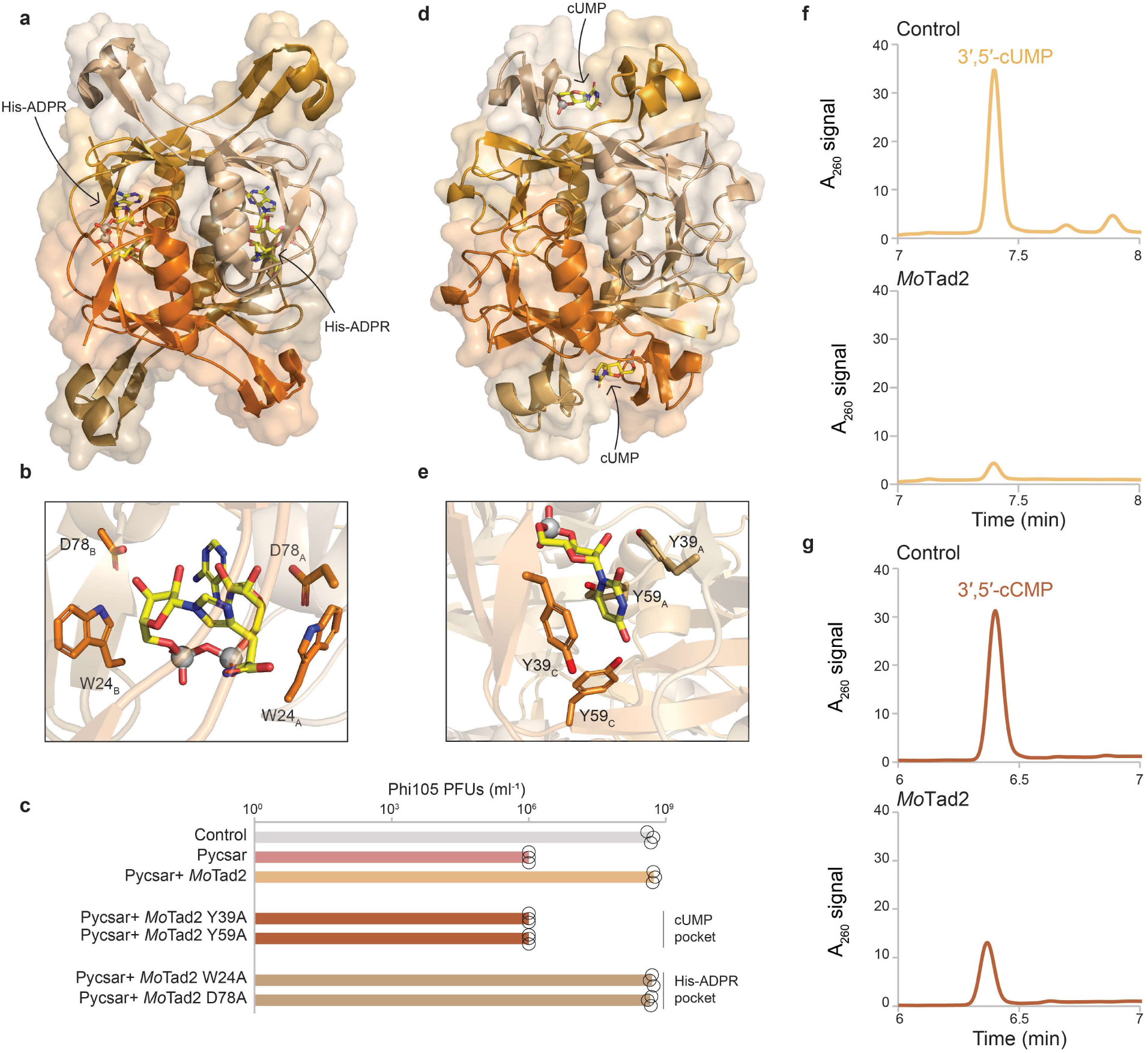
Molecular basis for suppression of Pycsar by *Mo*Tad2. (A) *Mo*Tad2 structure with the type II Thoeris ligand, His-ADPR (PDB: 9EIB)^5^. (B) Close-up view of *Mo*Tad2 His-ADPR binding pocket, highlighting key interacting residues. (C) Effect of point mutations in *Mo*Tad2 on the ability of the sponge to suppress the defensive activity of *Bacillus* sp. G1 Pycsar. Data represent PFUs per milliliter of phi105 phage infecting negative control cells (no system, grey), cells expressing Pycsar (red), or cells co-expressing Pycsar and a *Mo*Tad2 variant (shades of orange). Bars show the average of three replicates with individual data points overlaid. (D) AlphaFold3-predicted structure of *Mo*Tad2 with the Pycsar ligand 3′,5′-cUMP (ipTM= 0.91). (E) Close-up view of *Mo*Tad2 cUMP binding pocket, highlighting predicted key interacting residues. (F–G) HPLC analysis of 3′,5′-cUMP (F) or 3′,5′-cCMP (G) incubated with either purified GFP (control, top lane) or with purified *Mo*Tad2 at a 1:4 molar ratio.

We next purified *Mo*Tad2 and incubated it with cUMP and cCMP in vitro. The purified protein was able to sequester both molecules from solution, with a higher observed efficiency for binding cUMP as compared to cCMP (Fig. 4F,G).Purified *Mo*Tad2 was unable to sequester 2′,3′-cUMP, again demonstrating that this sponge is specific to the immune signaling molecule produced by Pycsar (Fig. S2D). Combined, our results describe the first phage sponge that inhibits Pycsar immunity.

### Inhibition of type IV Thoeris by Tad2 proteins

Type IV Thoeris is a defense system comprising a TIR-domain protein and a caspase-like protease. The TIR protein produces the signaling molecule N7-cADPR, an isomer of cADPR in which the ribose is bound to a nitrogen atom in position 7 in the adenine ring^4^. Once produced in response to phage infection, N7-cADPR binds a dedicated pocket in the caspase-like protease and activates it to indiscriminately degrade both phage and host proteins^4^. To date, no phage protein was known to inhibit type IV Thoeris.

Two Tad2 homologs in our set were found to inhibit the activity of type IV Thoeris from *Pseudomonas* sp. 1-7 when co-expressed with this system in *E. coli* K-12 MG1655 (Fig. 5A-C). To further verify the inhibitory function, we integrated one of these homologs (*tad2_71*) into the genome of phage T6, replacing the *acb1* gene naturally encoded in this phage^31^. While the wild type phage T6 was blocked with type IV Thoeris, phage T6 engineered to carry *tad2_71* was able to overcome type IV Thoeris defense (Fig. 5D).

**Figure 5.**
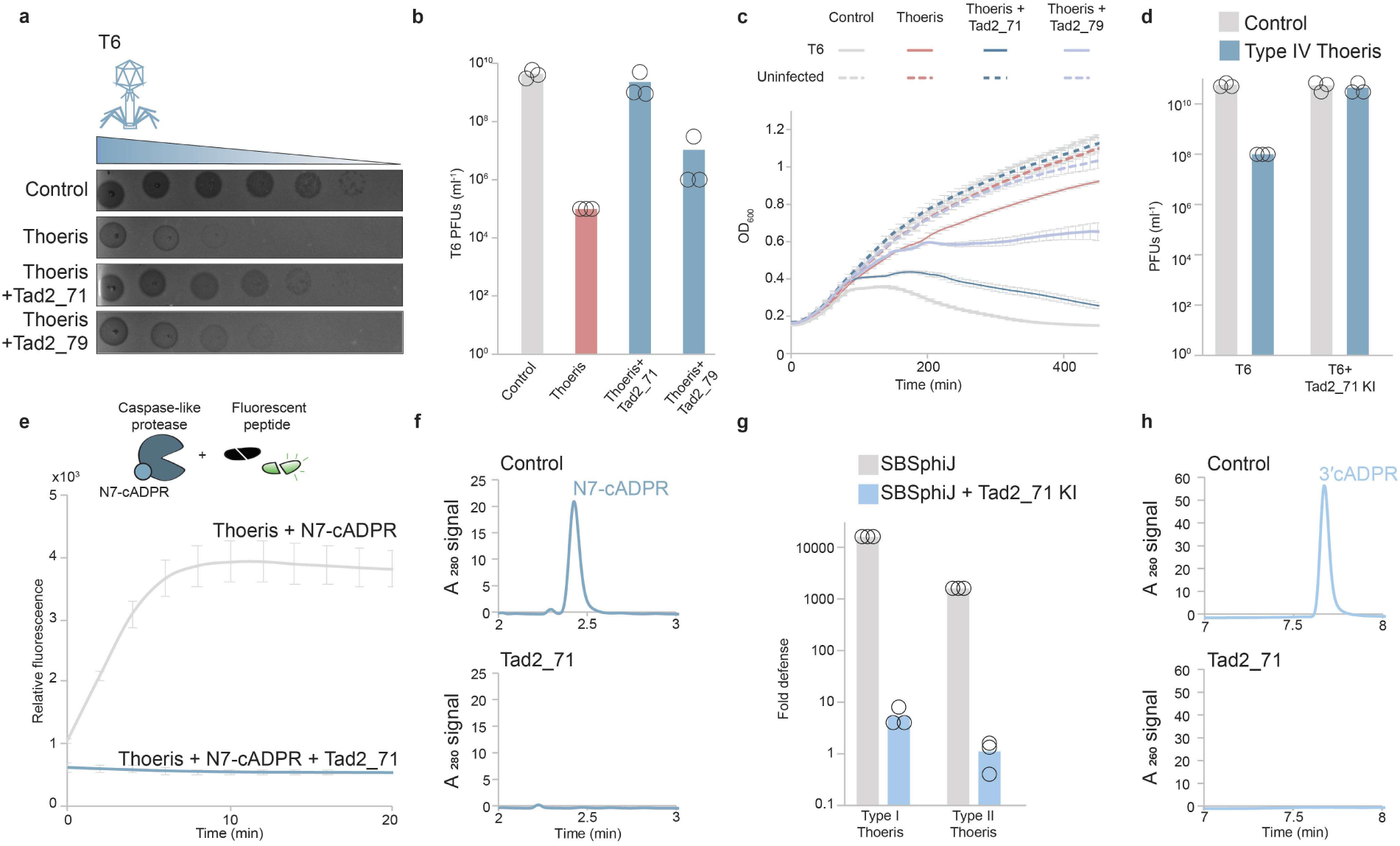
Tad2 homologs inhibit Type IV Thoeris. (A) Two Tad2 homologs capable of overcoming type IV Thoeris defense. Shown are tenfold serial dilution plaque assays, comparing the plating efficiency of phage T6 on bacteria that express the type IV Thoeris alone or with the Tad2 homolog. Images are representative of three replicates. (B) Plating efficiency of phage T6 on negative control *E. coli* cells (no system, grey), cells expressing type IV Thoeris (red), and cells co-expressing both type IV Thoeris and a homolog of Tad2 (blue). Data represent PFUs per milliliter, and bars show the average of three replicates with individual data points overlaid. (C) Growth curves of *E. coli* cells expressing type IV Thoeris (red), co-expressing type IV Thoeris and a Tad2 homolog (blue), or control cells expressing RFP and GFP instead (grey). Cells were infected with phage T6 at an MOI of 0.01. Each curve is the average of three replicates, with error bars indicating standard deviation. (D) A T6 phage engineered to express Tad2_71 overcomes type IV Thoeris defense. Data represent PFUs per milliliter of phages infecting *E. coli* cells expressing the type IV Thoeris system (blue) or control cells without the defense system (grey). Bars show the average of three biological replicates with individual data points overlaid. (E) Protease reporter assay showing the activity of the caspase-like protease in response to N7-cADPR pre-incubated either with purified GFP (grey) or Tad2_71 (blue). Curves represent the average of three replicates, error bars indicate standard deviation. (F) HPLC analysis of N7-cADPR after incubation with either purified GFP (control, top lane) or purified Tad2_71 at a 1:2 molar ratio. (G) A SBSphiJ phage engineered to express Tad2_71 overcomes type I and type II Thoeris defense in *B. subtilis*. Fold defense is calculated as the ratio between the plating efficiency on control cells that do not carry a defense system and cells expressing type I or II Thoeris. Data represent the average of three biological replicates with individual data points overlaid. (H) HPLC analysis of 3′cADPR incubated either with purified GFP (control, top lane) or Tad2_71 at a 1:2 molar ratio.

It was previously shown that when incubated with N7-cADPR, the protease from *Pseudomonas* sp. 1-7 type IV Thoeris efficiently cleaves the synthetic peptide EEKAR-7-amino-4-methylcoumarin (AMC) and releases a fluorescent signal^4^. We used this fluorescence-based protease activation assay to evaluate Tad2_71 activity in vitro. Our data show that when a solution containing N7-cADPR is pre-incubated with purified Tad2_71, it can no longer trigger protease activity, suggesting that Tad2_71 eliminates the signaling molecule (Fig. 5E). HPLC analysis verified that following incubation with Tad2_71, N7-cADPR could not be detected in solution (Fig. 5F).

Remarkably, data from our screens suggested that Tad2_71 can inhibit, in addition to type IV Thoeris, also types I and II Thoeris (Fig. 3A; Table S2). To test if this protein can inhibit types I and II Thoeris also when expressed from a phage genome, we engineered *tad2_71* into the genome of SBSphiJ, a *Bacillus* phage natively blocked by both type I and type II Thoeris. The engineered phage was able to overcome both type I and type II Thoeris, demonstrating that Tad2_71 is a general Thoeris inhibitor capable of hampering defense by the three types of Thoeris systems characterized to date (Fig. 5G). Indeed, HPLC analysis showed that Tad2_71 efficiently binds 3′cADPR as well (Fig 5H). We were unable to test Tad2_71 binding to His-ADPR in vitro, as this molecule could not be produced in large quantities, and is currently not commercially available.

## Discussion

In this study we report extensive functional diversity among phage sponges that sequester host immune signaling molecules. We documented the first sponges to inhibit Pycsar and type IV Thoeris, and showed that Acb2 homologs, previously considered as exclusively targeting CBASS-produced molecules, can also sequester 3′cADPR and inhibit Thoeris defense. We also verified previous observations that Tad1 sponges can inhibit both CBASS and Thoeris defense, and found that this activity is widespread among diverse homologs (Fig. S3)^25^.

Our data show that sponge proteins have high specificity to the molecules they bind: Acb2_43 binds 3′cADPR but not 3′3′-cGAMP, and *Mo*Tad2 inhibits type II Thoeris and Pycsar but not type I Thoeris or CBASS. The observed high specificity in molecule binding may be explained by recent discoveries on bacterial defense systems that produce “decoy” signaling molecules, generated as a bait for phage sponges^7,8^. These molecules are being constitutively produced in the bacterial cell regardless of phage infection, and serve to inhibit an otherwise toxic transmembrane-spanning protein. Sequestration of the decoy molecules by a phage sponge releases this inhibition and induces cellular toxicity that halts the phage infection process, so in these cases the sponge triggers bacterial immunity rather than inhibiting it. The decoy molecules resemble true immune signaling molecules: the molecule 2′3′-c-di-AMP serves as a bait for T4 Acb2, which naturally binds 3′3′-cGAMP and other CBASS signaling molecules to inhibit defense^21,22,29^. Presumably, these decoy molecules exert evolutionary pressure on phage sponges to adopt high specificity to the molecules they target but not to decoy molecules. Nevertheless, our findings report sponges, such as Tad2_71, that exhibit a broad range of binding specificities. This range can be beneficial in the phage–bacteria arms race, enabling phages to use a single small protein to counteract multiple types of bacterial immunity.

The topologies of the phylogenetic trees we constructed might suggest that specification of sponges towards binding new molecules may have evolved multiple times in the evolutionary history of a given sponge family. For example, while most tested Acb2 homologs inhibit CBASS defense, there seem to be four distinct clades on the Acb2 tree that contain sponges inhibiting Thoeris (Fig. 1A). Data from these trees, however, should be interpreted with caution. As the sponge proteins are short and show high sequence divergence, the positioning of many of the clades withing the phylogenetic analyses are unreliable, as measured by low bootstrap values (Fig. S4). Therefore, in the current study we used these trees as guidelines to select sponges that span the diversity of the family for our experiments, but do not refer to the trees as accurately reconstituting the evolutionary history of the studied sponge families.

Discoveries of defense systems that rely on small molecule signaling have accumulated rapidly in the past few years^1–8^, and it is estimated that many such systems remain to be discovered^12^. Sponge proteins proved useful for the discovery and characterization of new immune signaling molecules. For example, crystallization of Tad1 with molecules derived from TIR-domain proteins was crucial for the discovery of 3′cADPR and 2′cADPR as immune signaling molecules in bacteria and plants, respectively^19^; and *Mo*Tad2 supported the discovery of His-ADPR as the signaling molecule produced by type II Thoeris^5^. Notably, about one quarter of the sponge homologs we experimented with (19 of 84) did not inhibit any of the seven defense systems we examined (Table S2). While some of these proteins might specifically sequester molecules of CBASS variants not tested in our hands, others might bind molecules produced by defense systems that have not been discovered yet. Given the central roles that sponges played in past discoveries of signaling defense systems, we predict that the large set of sponges we collected will help in characterizing new immune signaling molecules in the future.

## Methods

### Identification of sponge proteins

∼32 million non redundant phage proteins from ∼2 million phage genome scaffolds were clustered based on sequence homology as previously described^26^, and ∼65,000 clusters of short proteins of unknown function with at least 20 members were further analyzed^26^. To generate multiple sequence alignment for an AlphaFold2 run, the databases of AlphaFold multimer 2.3.1^32^ with default parameters, with the IMG/VR v3 database^27^ added to the default protein databases, were used. AlphaFold multimer was then used to predict the structure of a representative sequence^33^ from each cluster folded as a dimer, trimer, tetramer, pentamer and hexamer, with a single model predicted (model 1) for each homo-oligomer. For predicted structures with a model confidence score^32^ of at least 0.8, five models were calculated, and structures with an average model confidence score of at least 0.8 over the five models were retained for further analysis. For proteins with more than one possible oligomeric state (score > 0.8), the highest oligomeric form was chosen for further analysis.

To search for clusters corresponding to known sponges, Clustal omega^34^ version 1.2.4 was used with default parameters to generate a multiple sequence alignment (MSA) for each cluster. Then, HHpred^35,36^ with default parameters was run for each MSA, using the databases pfam35^37,38^ and PDB70^39^ downloaded on October 24, 2023. All proteins belonging to clusters annotated (with HHpred probability higher than 0.9) as Tad1 (7UAW), Tad2 (PF11195), and Acb2 (7UQ2 or 8H39) were collected for downstream analysis.

Next, each protein sequence from the selected clusters was further analyzed individually using HHblits and HHsearch^40^ with default parameters using databases pfam35^37,38^ and PDB70^39^ downloaded on October 24, 2023. Proteins not showing homology to the respective sponge when analyzed individually were removed from the analysis. Proteins longer than 500 amino acids were removed, and proteins containing the domains PF21825 or PDB 6E3C in addition to the sponge domain were also removed. Previously studied sponge homologs were added manually to the protein datasets. Overall, 2953 sequences (1587 non redundant sequences) were identified as Tad1 homologs, 7492 (2783 non redundant) as Tad2, and 9141 (4204 non redundant) as Acb2 (Table S1).

The protein sequences were aligned using MAFFT version 7.520^41^. The phylogenetic tree was constructed using IQ-TREE v2.2.0^42^ with the parameters -m LG and -B 1000, and visualized by the iTOL web application (v5)^43^.

### Bacterial strains and growth conditions

*E. coli* K-12 MG1655 and *B. subtilis* BEST7003 strains were grown in magnesium manganese broth (MMB; LB + 0.1 mM MnCl_2_ + 5 mM MgCl_2_) at 37 °C or 30 °C with shaking at 200 RPM. Whenever applicable, the appropriate antibiotics were added at the following concentrations: For *B. subtilis* strains spectinomycin (100 μg ml^−1^) and chloramphenicol (5 μg ml^−1^), and for *E. coli* strains spectinomycin (50 μg ml^−1^) and kanamycin (50 μg ml^−1^).

Type I Thoeris from *Bacillus cereus* MSX-D12 (producing 3′cADPR), type II Thoeris from *Bacillus amyloliquefaciens* Y2 (His-ADPR), were previously cloned under their native promoters into the *amy*E locus of the *B. subtilis* BEST7003 genome^44^. Pycsar from *Bacillus* sp. G1 (3′,5′-cUMP) was cloned similarly under its native promoter to the *amyE* locus. An empty vector was integrated as a negative control.

Type I CBASS from *E. albertii* MOD1-EC1698 (3′3′-cGAMP)^28^, Type I CBASS from *Y. aleksiciae* IP28587 (3′,3′-cUA)^28^, type IV Thoeris from *Pseudomonas* sp. 1-7 (N7-cADPR)^4^, Pycsar from *E. coli* E831 (3′,5′-cCMP)^3^ and RFP as negative control were each integrated into the genome of *E. coli* MG1655 using the Tn7 integration plasmid (downstream of the gmlS gene, spectinomycin resistance, induced by anhydrotetracycline, Sigma cat.37919)^45^.

Plasmids were synthesized by Twist Bioscience or GenScript Corporation unless stated otherwise. Transformation into *E. coli* K-12 MG1655 using standard electroporation or TSS, or in *B. subtilis* using MC medium, was performed as previously described^44,46^

### Phage strains

The *B. subtilis* phage SBSphiJ (GenBank: LT960608.1) was previously isolated by us and was used for all infection experiments involving type I and type II Thoeris. The *B. subtilis* phage Phi105 (GenBank: NC_048631.1) was obtained from the DSMZ (DSM HER46) and used to infect the Pycsar system from *Bacillus* sp. G1. The *E. coli* phages T2 (DSMZ 16352) and BAS60 (from the BASEL phage collection, kindly provided by Prof. Alexander Harms)^47^ were both used to infect type I CBASS from *E. albertii*. The *E. coli* phage T6 (DSMZ 4622) was used to infect type IV Thoeris from *Pseudomonas* sp. 1-7. The *E. coli* phages T5 and P1, kindly provided by Udi Qimron, were used to infect Pycsar from *E. coli* E831 (both phages) and CBASS from *Y. aleksiciae* IP28587 (T5 only).

The phages were propagated on either *E. coli* MG1655 or *B. subtilis* BEST7003 by picking a single phage plaque into a liquid culture grown at 37 °C to an optical density at 600 nm (OD_600_) of 0.3 in MMB broth until culture collapse (or 3 hours in the case of no lysis). The culture was then centrifuged for 10 min at 3,200 *g* and the supernatant was filtered through a 0.2 µm filter to get rid of remaining bacteria and bacterial debris.

### Cloning and transformation of sponge proteins

Acb2, Tad1 and Tad2 homolog genes and GFP as a control were codon optimized for *E. coli* and synthesized by Genscript Corp. under the control of IPTG promoter. These constructs were subsequently cloned into the thrC-Phspank^19^ vector, which carries a low-copy pSC101 origin of replication and a kanamycin resistance marker. The resulting plasmid was then introduced into *B. subtilis* BEST7003 cells, where the respective defense system was integrated into the *amy*E locus, or into *E. coli* K-12 MG1655, where the defense system was integrated downstream of the gmlS gene and induced by anhydrotetracycline.

Transformations were first performed using a 96-well format. *B. subtilis* BEST7003 cells containing the defense system were cultured in MC medium at 37 °C for 3 hours, using the protocol described by Doron et al^44^. MC medium was composed of 80 mM K2HPO4, 30 mM KH2PO4, 2% glucose, 30 mM trisodium citrate, 22 μg ml^−1^ ferric ammonium citrate, 0.1% casein hydrolysate (CAA) and 0.2% potassium glutamate. Aliquots of 200 µL were transferred to a deep 96-well plate, and 200 ng of the plasmid with the sponge gene or control sequence was added to each well. After 3 hours incubation at 37 °C, 1 mL of MMB medium supplemented with spectinomycin (100 μg/mL) and chloramphenicol (5 μg/mL) was added to each well to dilute the MC medium, and the plate was incubated overnight at 30 °C. On the following day, a 10 µL aliquot from each well was transferred into 990 µL of fresh MMB containing spectinomycin (100 μg/mL) and chloramphenicol (5 μg/mL) for a single passage. After overnight incubation at 37 °C, the cultures were used as starters for infection experiments in liquid cultures.

For *E. coli* K-12 MG1655, cells carrying the defense system integrated into the genome were grown in MMB medium until reaching an OD_600_ of 0.3. The cells were then centrifuged at 3,200 *g* for 5 minutes and resuspended in TSS medium to a final cell concentration of 20X^20^. A 50 µL aliquot of the cell suspension was distributed into a deep 96-well plate, and 50 ng of the plasmid with the anti-defense candidate or control sequence (GFP) was added to each well. The transformation protocol consisted of three incubation steps: 5 minutes on ice, 5 minutes at room temperature, and an additional 5 minutes on ice. Subsequently, 500 µL of antibiotic-free MMB was added to each well, followed by incubation with shaking at 37 °C for 1 hour. After this recovery period, 500 µL of MMB containing spectinomycin (100 μg/mL) and kanamycin (100 μg/mL) was added to each well, to reach a final concentration of 50 μg/mL spectinomycin and 50 μg/mL kanamycin, and the plate was incubated overnight at 37 °C. On the next day, 10 µL from each well was transferred into 990 µL of fresh MMB containing spectinomycin (50 μg/mL) and kanamycin (50 μg/mL) for a single passage. After overnight incubation at 37 °C, the cultures were used as starters for infection experiments in liquid cultures. (Table S2).

### Determination of anti-defense phenotypes

Cells transformed in 96-well plates were infected in liquid culture with phages typically restricted by the corresponding defense system. The phages used for each defense system are detailed in the “Phage strains” section. Genes that either failed to transform or showed a potential anti-defense phenotype in this format were reintroduced individually (as described in the “Bacterial strains and growth conditions” section) into cells carrying the relevant defense system (Table S2). These strains were subsequently challenged with the corresponding phage in plaque assays. A sponge homolog was defined as having an anti-defense phenotype if, in plaque assays, the average of three replicates showed at least a 10-fold reduction in defense compared to the system-only control. (Table S2).

### Infection in liquid culture

Overnight cultures of bacteria harboring the defense system, negative control lacking the system and bacteria encoding the system and a sponge gene were diluted 1:100 in MMB medium. For both *B. subtilis* and *E. coli* cultures, 1 mM IPTG was added to induce expression of candidate anti-defense proteins. For *E. coli* cultures anhydrotetracycline (aTc, Sigma cat.37919) was added to induce the expression of the defense system: (800 nM aTc for *E. albertii* CBASS, 400 nM for *Y. aleksiciae* CBASS, *Pseudomonas* sp. Thoeris and *E. coli* Pycsar and 40 nM aTc in the experiment with T5 *Mo*Tad2 KI and *E. coli* Pycsar). For *B. subtilis* cultures, expression was driven from the native promoter of the respective defense system. Cells were incubated at 37 °C while shaking at 200 rpm until early log phase (OD_600_ of 0.3) and infected in a 96-well plate containing 20 μL of phage lysate. Plates were incubated at 30 °C or 37 °C with shaking in a TECAN Infinite200 plate reader and OD_600_ was followed with measurement every 10 min.

### Plaque assays

Phage titer was determined using the small drop plaque assay method^48^. An overnight culture (300 µL) of *E. coli* or *B. subtilis* was mixed with 30 mL MMB 0.5% agar supplemented with 1mM IPTG. The agar was also supplemented with aTc as follows: 800 nM aTc for *E. albertii* CBASS, 400 nM aTc for *Y. aleksiciae* CBASS and *E. coli* Pycsar, 400 nM aTc for experiments with *Pseudomonas* sp. type IV Thoeris unless otherwise stated, and 50 nM aTc in the experiment with T6 Tad2_71 KI and *Pseudomonas* sp. type IV Thoeris. Agar was poured into a 10-cm square plate followed by incubation for 1 h at room temperature. Tenfold serial dilutions in MMB were carried out for each of the tested phages and 10 µl drops were put on the bacterial layer. After the drops had dried up, the plates were inverted and incubated overnight at room temperature (for type II Thoeris and *Y. aleksiciae* CBASS) or 30 °C (for type I Thoeris and *Bacillus* sp. Pycsar) or 37 °C (for type IV Thoeris, *E. albertii* CBASS and *E. coli* Pycsar). PFUs were determined by counting the derived plaques after overnight incubation and lysate titer was determined by calculating PFUs per milliliter. When no individual plaques could be identified, a faint lysis zone across the drop area was considered to be 10 plaques. The efficiency of plating was measured by comparing plaque assay results for control bacteria and those for bacteria containing the defense system and/or the defense system with a candidate anti-defense gene.

### Generation of phage knock-in

*<u>Phage SBSphiJ with Acb2_43:</u>* The DNA sequence of *acb2_43* was amplified from the respective Genscript-synthesized pSG-thrC-Phspank plasmid using KAPA HiFi HotStart ReadyMix (Roche, cat. #KK2601) with the following primer pair:

F: ttttccctccagttgttcttatgcgtttccttcaccgtatttg

R: gagacctaccgactgagtaacgagggtaaatgtgagcactcac

The backbone fragment, containing ∼1.2 kb upstream and downstream genomic arms for integration, was amplified from the plasmid used previously for knock-in of the *tad1* gene into SBSphiJ^19^ using the primer pair F: gaacaactggagggaaaacacttg, R: agctacctcctataggtataaaatttg.

Cloning was performed using the NEBuilder HiFi DNA Assembly kit (NEB, cat. #E5520S), and the assembled vector was transformed into NEB 5-alpha competent cells. The final plasmid was integrated into the *thr*C locus of *B. subtilis* BEST7003. The *acb2_43*-containing *B. subtilis* BEST7003 strain was then infected with phage SBSphiJ at a multiplicity of infection (MOI) of 0.1, and the resulting lysate was collected. Phages containing the knock-in were selected using a Cas13a-gRNA-SBSphiJ strain^49^ as previously described^20^on agar plates with 0.2% xylose to induce Cas13 expression. Resistant phages were purified three times on *B. subtilis* BEST7003. Integration of the anti-defense candidate was confirmed by PCR and sequencing.

*<u>Phage T5 with MoTad2</u>*: The T5 knock-in was generated using homologous recombination and Cas13a-based selection. A gRNA targeting the upstream region of *gene A1* cloned into the plasmid pBA559^49^ using the following primer pair:

F: cgttataatttgcaacattaatttaatgcttgggcccgaacaaaaac

R: attaatgttgcaaattataacgcctatgttttagtccccttcatttttggggt

Cloning was performed with the NEBuilder HiFi DNA Assembly kit (NEB, cat. #E5520S), and the vector was introduced into NEB 5-alpha competent cells. Clones were validated by plasmid sequencing (Plasmidsaurus), and the plasmid was transformed into *E. coli* MG1655. A homology recombination template was prepared by cloning the amplified coding sequence of *Mo*Tad2 from the respective Genscript-synthesized thrC-Phspank template, including the upstream ribosome binding site, into the pUC19 plasmid (digested with SmaI endonuclease) using the following primer pair:

F: taacggtttgtttttctgcggaaataaccataataataaaactccattaaagggattaaaaaacgtttgattgccgaccttgactagtgc

R: ctgatttgttttattgccgccgttgcttgtggtgttagttctagccttatgcctttaaattaatgttagtcgacagctagctgattaactaataag.

The vector was transformed into NEB 5-alpha cells, validated by plasmid sequencing (Plasmidsaurus), and then transformed into *E. coli* MG1655 for recombination and knock-in selection (described below in the “Phage recombination and Cas13a-based” subsection).

*<u>Phage T6 with Tad2_71</u>*: The T6 knock-in was similarly generated by homologous recombination and Cas13a-based selection. A gRNA targeting the coding region of the *acb1* gene in phage T6 was cloned into the pBA559 plasmid (Addgene #186235)^49^ using the following primer pair:

F: aaagacccgttgaaaagtctttaaaatgcttgggcccgaacaaaaac

R: aaagacttttcaacgggtctttatgtagttttagtccccttcatttttggggt

The gRNA plasmid was assembled using the NEBuilder HiFi DNA Assembly kit and transformed into NEB 5-alpha competent cells. Correct clones were verified by plasmid sequencing (Plasmidsaurus) and transformed into *E. coli* MG1655. A homology recombination template was generated by amplifying the Tad2_71 coding sequence, with a synthetic ribosome binding site, from the respective Genscript-synthesized thrC-Phspank template using the following primer pair:

F: tatttacttcctcggtattataacaccatagctacaggaggataataaaatggaaaataatgtcatcgagactgtagattttggc

R: gcatgtaaacaactttgtgaaactacttagatttaatgatccaatcctcgg.

To amplify the plasmid backbone, pGEM9z was used with primer pair:

F: tttcacaaagttgtttacatgctgatgaggtagtgatactattatctcatcaagaattcgtcgacgagctccc

R: ttataataccgaggaagtaaataactttttatcgttttgttattcttcatcttttgcttcatcagtagattcagcagtaagattcttgacagcttactagtgatgcatattctatagtgtcacc.

Cloning was performed with the NEBuilder HiFi DNA Assembly kit, transformed into NEB 5-alpha cells, validated by plasmid sequencing (Plasmidsaurus), and transformed into *E. coli* MG1655 for recombination and knock-in selection (see below).

*<u>Phage recombination and Cas13a-based selection (T5 and T6):</u> E.* coli MG1655 strains carrying the appropriate recombination plasmid were infected with phage T5 or T6 at MOI 0.01 in 5 mL MMB medium. After 3 hours of incubation at 37 °C, the lysate was harvested, filtered through a 0.4 µm filter, and stored at 4 °C. Cas13a-based selection was performed by mixing 100 µL of overnight *E. coli* cultures expressing Cas13a and the relevant gRNA with 15 mL of 0.5% MMB top agar supplemented with 10 nM anhydrotetracycline. The mixture was poured onto round Petri dishes and allowed to dry at room temperature for 1 hour. 1 µL of filtered phage lysate was spotted onto the plates, which were incubated overnight at 37 °C. Plaques were collected and screened by PCR and sequencing.

### Preparation of filtered cell lysates for NADase Enzymatic assay

For generating filtered cell lysates, *B. subtilis* BEST7003 cells co-expressing the sponge homolog and the *B. cereus* MSX-D12 Thoeris system in which ThsA was inactivated (ThsB + ThsAN112A) (as described previously)^2^ were used. The candidates (Acb2_11, Acb2_31, and Acb2_43) were integrated in the thrC locus as described above and expressed from an inducible Phspank promoter. Controls included cells expressing only the ThsB + ThsA(N112A) Thoeris system with GFP (negative control) or with SBSphiJ7 Tad1 (positive control). All cultures were grown overnight and then diluted 1:100 in 250 mL MMB medium supplemented with 1 mM IPTG and grown at 37 °C, 200 rpm shaking for 120 min followed by incubation and shaking at 25 °C, 200 rpm until reaching an OD_600_ of 0.3. At this point, a sample of 50 mL was taken as the uninfected sample (time: 0 min), and the SBSphiJ phage was added to the remaining culture at an MOI of 10. Flasks were incubated at 25 °C with shaking (200 rpm) for the duration of the experiment. Samples of 50 mL were collected 105 minutes post infection. Immediately after sample removal the sample tubes were centrifuged at 4 °C for 10 min at 3,200 *g* to pellet the cells. The supernatant was discarded, and the pellet was flash frozen and stored at −80 °C. To extract cell metabolites from frozen pellets, 600 μL of 100 mM Na phosphate buffer (pH 8.0) was added to each pellet. Samples were transferred to FastPrep Lysing Matrix B in a 2-mL tube (MP Biomedicals, cat #116911100) and lysed at 4 °C using a FastPrep bead beater for 2 × 40 s at 6 ms^−1^. Tubes were then centrifuged at 4 °C for 10 min at 15,000 *g*. The supernatant was then transferred to an Amicon Ultra-0.5 Centrifugal Filter Unit 3 kDa (Merck Millipore, no. UFC500396) and centrifuged for 45 min at 4 °C, 12,000 *g*. Filtered lysates were taken for in vitro ThsA-based NADase activity assay.

### NADase Enzymatic assay

The ThsA protein was expressed and purified as previously described^20^, and ThsA-based NADase activity assay for the detection of 3′cADPR was carried out, as previously described^2^. NADase reaction was carried out in black 96-well half-area plates (Corning, cat #3694). In each reaction microwell, purified ThsA protein was added to cell lysate, or to 100 mM sodium phosphate buffer pH 8.0. A 5 μL volume of 5 mM nicotinamide 1,N6-ethenoadenine dinucleotide (εNAD+, Sigma, cat #N2630). Solution was added to each well immediately before the beginning of measurements, resulting in a final concentration of 100 nM ThsA protein in a 50 µL final volume reaction. Plates were incubated inside a Tecan Infinite M200 plate reader at 25 °C, and measurements were taken at 300 nm excitation wavelength and 410 nm emission wavelength. The reaction rate was calculated from the linear part of the initial reaction.

### Protein expression and purification for biochemical assays

GFP, Tad2_71, and Acb2_43 were cloned into the pET28-His-bdSUMO expression vector (TWIST Bioscience, USA), which encodes an N-terminal His14-bdSUMO tag. This vector was constructed by transferring the His14-bdSUMO cassette from the K151 plasmid (generously provided by Prof. Dirk Görlich, Max Planck Institute, Göttingen, Germany) into the pET28-TevH backbone^50,51^. Constructs were transformed into *E. coli* LOBSTR-BL21(DE3)-RIL cells (Kerafast, USA), and cultures were grown in MMB medium (1 L, supplemented with 100 µg/mL kanamycin) at 37 °C with shaking at 200 rpm. At OD_600_ ≈ 0.6, protein expression was induced with 1 mM IPTG, followed by overnight incubation at 16 °C with shaking.

Cells were harvested by centrifugation (20,000 *g*, 30 min, 4 °C), and pellets were either processed immediately or stored at −80 °C. Lysis was performed on ice using an EmulsiFlex-C3 homogenizer (Avestin, Canada) in lysis buffer. For GFP and Tad2_71, the buffer contained 20 mM HEPES pH 7.3, 0.4 M NaCl, 30 mM imidazole, and 1 mM DTT; for Acb2_43, Tris-HCl pH 7.5 was used in place of HEPES. After clarification (20,000 *g*, 20 min, 4 °C) and filtration (0.4 µm for GFP/Tad2_71; 0.2 µm for Acb2_43), lysates were applied to a 5 mL HisTrap FF column (Cytiva), pre-equilibrated with lysis buffer. The column was washed with lysis buffer, followed by a high-salt wash (1 M NaCl), and the protein was eluted with a buffer containing 300 mM imidazole, 0.4 M NaCl, 1 mM DTT, and either 20 mM HEPES pH 7.3 (GFP/Tad2_71) or 20 mM Tris-HCl pH 7.5 (Acb2_43).

Eluted proteins (∼15 mL) were dialyzed overnight at 4 °C in 3.5 kDa MWCO Slide-A-Lyzer™ cassettes (Thermo Scientific) in cleavage buffer (20 mM HEPES for GFP and Tad2_71 or Tris-HCl for Acb2_43, 125–150 mM KCl, 1 mM DTT), with the addition of 1 µM His-tagged bdSENP1 protease. After cleavage, samples were centrifuged and supplemented with NaCl and imidazole to final concentrations of 275 mM and 40 mM, respectively. Samples were reloaded onto a HisTrap FF column to remove uncleaved protein, the bdSUMO tag, and the protease. The flowthrough, containing the cleaved, untagged protein, was collected. For Acb2_43, the flowthrough was additionally washed three times with 20 mM Tris-HCl pH 7.5, 150 mM NaCl using a 10 kDa Vivaspin centrifugal concentrator (Cytiva). All proteins were stored at −80 °C.

The *Mo*Tad2 protein used in this study was expressed and purified as previously described^5^

### In vitro sponge-ligand experiments

Sponges were expressed and purified as described above in the “protein expression and purification” section. 80 µM Acb2_43 with 20 µM 3′cADPR (BioLog cat. C 404), 40 µM Tad2_71 with 20 µM 3′cADPR or N7-cADPR^4^ or 20 µM *Mo*Tad2 with 5 µM 3′,5-cCMP, 3′,5′-cUMP or 2′,3′-cUMP (BioLog Cat# C 001, U 001, U004) were mixed in 100 mM sodium phosphate buffer pH 7.2-7.6. Ligands were incubated with sponge proteins or with purified GFP as control for 10 minutes at room temperature. For GFP, buffer was adjusted to match the corresponding sponge protein. Following the incubation the samples were filtered through a 3 kDa MWCO filter, and flow through was collected and analyzed by HPLC.

20 µL of the samples were analyzed using HPLC. HPLC of the obtained fraction was performed using Agilent 1260 and chromatography SUPELCOSIL™ LC-18-T HPLC Column. The following protocol was used for all runs: 1 min of mobile phase A 100%, 2 min 75% A and 25% B, 2 min 50% A and 50% B, 2 min 20% A and 80% B and 3 min 100 % A, 1 mL/min flow rate. Mobile phase A was 20 mM potassium phosphate pH 6 and B was 20 mM potassium phosphate pH 6 in 20% methanol.

### Acb2_43 expression and purification for crystallization

Acb2_43 was expressed and purified from *E. coli*. A synthetic gene fragment (Integrated DNA Technologies) was cloned by Gibson assembly (NEB) into a custom pET vector containing a 6×His-tagged N-terminal human SUMO2 fusion. Plasmids were transformed into *E. coli* BL21(DE3)-RIL cells (Agilent) and plated on MDG media (1.5% Bacto agar, 0.5% glucose, 2 mM MgSO_4_, 0.25% aspartic acid, and trace metals) with appropriate antibiotic selection (100 µg mL^−1^ ampicillin and 34 µg mL^−1^ chloramphenicol). Transformed colonies were used to inoculate a ∼30 mL starter culture and grown overnight for ∼17 hours in MDG media at 37°C with shaking (230 rpm). Expression cultures were inoculated with ∼15 mL of MDG starter culture in 1 L of M9ZB media (47.8 mM Na_2_HPO_4_, 22 mM KH_2_PO_4_, 18.7 mM NH_4_Cl, 85.6 mM NaCl, 1% casamino acids, 0.5% glycerol, 2 mM MgSO_4_, trace metals, 100 µg mL^−1^ ampicillin, and 34 µg mL^−1^ chloramphenicol) and grown to an OD_600_ of ∼2.5-3.0 prior to induction with 0.5 mM isopropyl-β-D-thiogalactoside (IPTG) for ∼15 hours at 16°C with shaking (230 rpm). After overnight expression, cultures were harvested by centrifugation at 3,500 × g for 20 minutes. The cell pellets were resuspended in 60 mL of lysis buffer (20 mM Tris-HCl pH 7.5, 400 mM NaCl, 10% glycerol, 30 mM imidazole, and 1 mM DTT) and lysed by sonication to release soluble proteins. The lysate was clarified by centrifugation at 50,000 × g for 30 minutes and the resultant supernatant was purified by nickel affinity chromatography. Briefly, the lysate was poured over 8 mL of Ni-NTA Agarose (Qiagen) in a gravity flow column. The resin was washed sequentially with 20 mL of lysis buffer, 70 mL of wash buffer (20 mM Tris-HCl pH 7.5, 1 M NaCl, 10% glycerol, 30 mM imidazole, and 1 mM DTT) and 35 mL of lysis buffer. The protein was eluted with 20 mL of elution buffer (20 mM Tris-HCl pH 7.5, 400 mM NaCl, 10% glycerol, 300 mM imidazole, and 1 mM DTT). The eluate was treated with ∼200 µg of recombinant human SENP2 protease to facilitate cleavage of the 6×His-SUMO tag and dialyzed overnight at 4°C in 14 kDa molecular weight cutoff dialysis tubing (Ward’s Science). Acb2_43 was dialyzed into dialysis buffer (20 mM Tris-HCl pH 7.5, 250 mM KCl, and 1 mM TCEP), concentrated using a 10 kDa MWCO centrifugal filter (Millipore Sigma) then further purified by size-exclusion chromatography using a 16/600 Superdex 75 column (Cytiva).

### Acb2_43–3′cADPR crystallization and structure determination

Crystals of the Acb2_43–3′cADPR complex were grown using hanging-drop vapor diffusion method for ∼12 days at 18 °C. Recombinant Acb2_43 was diluted to 10 mg mL^−1^ in a buffer containing 20 mM Tris-HCl pH 7.5, 250 mM KCl, and 1 mM TCEP. The diluted protein was incubated with 500 μM 3′cADPR (Biolog Life Science Institute) on ice for ∼30 minutes. The resultant protein mixture was allowed to equilibrate to 18 °C and crystals were grown in 96-well trays containing 70 μL reservoir solution and 400 nL drops. Drops were mixed 1:1 with purified protein and reservoir solution (20% (w/v) PEG-1500 and 0.1 M HEPES pH 7.5). Crystals were cryo-protected with reservoir solution supplemented with 22% (v/v) ethylene glycol and 500 μM 3′cADPR (Biolog Life Science Institute) and harvested by flash-freezing in liquid nitrogen. X-ray diffraction data were collected at Advanced Photon Source (beamline 24-ID-E), and data were processed on the RAPDv2 platform using XDS^52^. Experimental phase information was determined by molecular replacement using Phaser-MR in PHENIX^53^ and a model of Acb2_43 generated using AlphaFold3^30^. Model building was performed using Coot^54^ and refined using PHENIX. A summary of crystallographic statistics is provided in Table S3. All structural figures were generated using PyMOL (Version 2.5.4, Schrödinger, LLC).

### Toxicity assay with Pycsar molecules

Medium-copy-number p15A origin of replication thrC-Phspank plasmid expressing *Mo*Tad2 or GFP as control^5^ were introduced to *E. coli* MG1655 carrying genomically-integrated *E. coli* E831 Pycsarfor toxicity assays. Overnight cultures of cells carrying Pycsar system from either *E. coli* E831 or *Bacillus* sp. G1 together with *Mo*Tad2, a negative control lacking the system, and a system-only control expressing GFP were diluted 1:100 in MMB medium. For both *B. subtilis* and *E. coli*, 1 mM IPTG was added to induce expression of the sponge. In *E. coli* cultures, 1000 nM anhydrotetracycline was also added to induce expression of the Pycsar system. Cells were incubated at 37 °C with shaking at 200 rpm until early log phase (OD_600_ ≈ 0.3). Then, cyclic nucleotide molecules or water as control were added in a 96-well plate to a final volume of 200 µL per well. For *E. coli*: 350 µM 3′,5′-cCMP (BioLog Cat# C 001) or 3′,5′-cUMP (BioLog Cat# U 001) were added. For *B. subtilis*: 500 µM 3′,5′-cUMP or 2′,3′-cUMP (BioLog Cat# U 004) were added. Plates were incubated at 37 °C (*E. coli*) or 30 °C (*B. subtilis*) with shaking in a TECAN Infinite200 plate reader, and OD_600_ was recorded every 10 minutes.

### Prediction of *Mo*Tad2 structure with cUMP

AlphaFold3^30^ with default parameters was used to generate structural models for *Mo*Tad2 with the ligand 3′,5′-cUMP. Four monomers of *Mo*Tad2 were used as an input together with two copies of 3′,5′-cUMP using the SMILES code: C1[C@@H]2[C@H]([C@H]([C@@H](O2)N3C=CC(=O)NC3=O)O)OP(=O)(O1)O.

### Type IV Thoeris protease reporter assays

Caspase-like protease, synthetic EEKAR-aminomethylcoumarin (AMC) conjugated peptides and N7-cADPR were produced as described previously^4^. Tad2_71 or control GFP (purified as described above) in final concentration of 0.4 µM were mixed with N7-cADRP in final concentration of 0.1 µM and incubated for 10 minutes at room temperature. The samples were mixed with 1 µM caspase-containing lysate^4^, 2 µM AMC peptides^4^ and 100 mM sodium phosphate buffer (pH 7.4) in a black 384-well plate. AMC fluorescence (excitation/emission wavelength: 341/441 nm) was measured on a Tecan Infinite M200 plate reader at 37 °C every 2 min and compared relative to a pure solution of 10 µM AMC.

## Acknowledgements

We thank members of the Sorek and Kranzusch labs for constructive criticism on the manuscript. R.S. was supported, in part, by the European Research Council (grant ERC-AdG GA 101018520), the Israel Science Foundation (MAPATS grant 2720/22), the Deutsche Forschungsgemeinschaft (SPP 2330, grant 464312965), the Minerva Foundation with funding from the Federal German Ministry for Education and Research, a research grant from the Estate of Hermine Miller, the Institute for Environmental Sustainability (IES) and the Center for Immunotherapy at the Weizmann Institute of Science, and the Knell Family Center for Microbiology. N. B. was supported by a postdoctoral grant from the Azrieli foundation.

I.O. was supported by the Ministry of Absorption New Immigrant program. E.Y. is supported by the Clore Scholars Program and, in part, by the Israeli Council for Higher Education (CHE) via the Weizmann Data Science Research Center. The research was supported in part by grants to P.J.K. from the Burroughs Wellcome Fund PATH program, The G. Harold and Leila Y. Mathers Charitable Foundation, the Cancer Research Institute, the Parker Institute for Cancer Immunotherapy, the Massachusetts Consortium on Pathogen Readiness (MassCPR), the Gates Foundation (INV-083469), and the National Institutes of Health (1DP2GM146250-01). R.B.C. is supported through a Landry Cancer Biology Research Fellowship (Harvard Faculty of Arts and Sciences). X-ray data were collected at the Northeastern Collaborative Access Team beamlines, which are funded by the National Institute of General Medical Sciences from the National Institutes of Health (P30 GM124165). The Eiger 16M detector on the 24-ID-E beam line is funded by a NIH-ORIP HEI grant (S10OD021527). This research used resources of the Advanced Photon Source, a U.S. Department of Energy (DOE) Office of Science User Facility operated for the DOE Office of Science by Argonne National Laboratory under Contract No. DE-AC02-06CH11357. This publication resulted from the data collected using the beamtime obtained through NECAT BAG proposal # 311950.

## Data Availability Statement

Coordinates and structure factors of Acb2 homolog 43 in complex with 3′cADPR have been deposited in the PDB under the accession code 9PTQ.

## Competing interests

R.S. is a scientific cofounder and advisor of BiomX and Ecophage. The other authors declare no competing interests.

## Supplementary Materials

**Figure S1.**
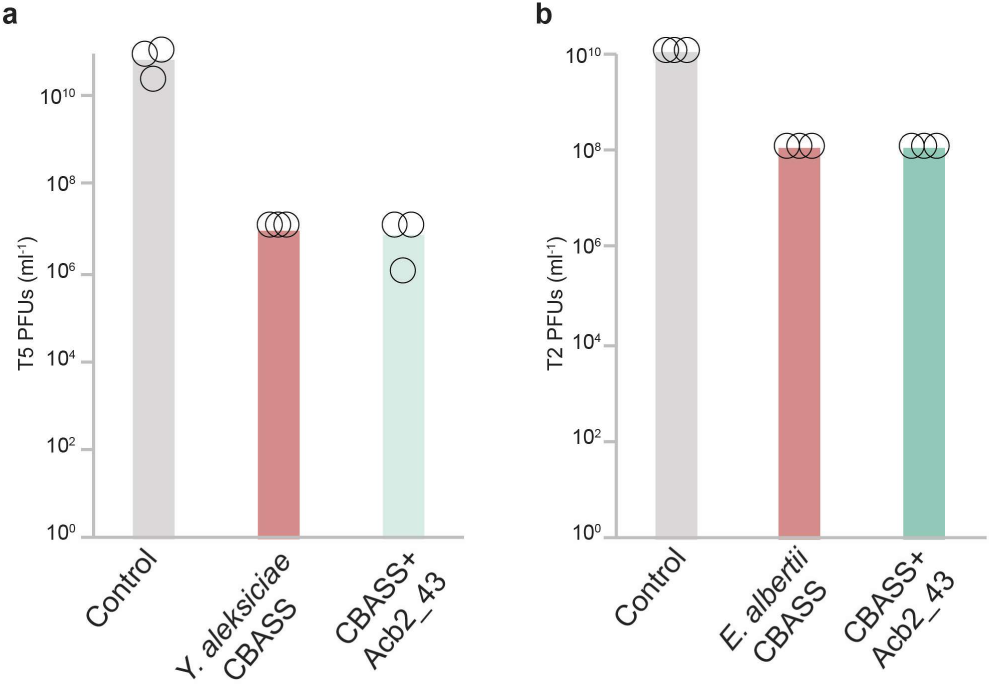
Acb2_43 does not affect CBASS defense. (A) Plating efficiency of phage T5 on negative control *E. coli* cells (grey), cells expressing the CBASS system from *Y. aleksiciae* (red), and cells co-expressing CBASS and Acb2_43 (green). (B) Plating efficiency of phage T2 on negative control *E. coli* cells (grey), cells expressing the CBASS system from *E. albertii* (red), and cells co-expressing CBASS and Acb2_43 (green). In both panels, data represent PFUs per milliliter, and bars show the average of three replicates with individual data points overlaid.

**Figure S2.**
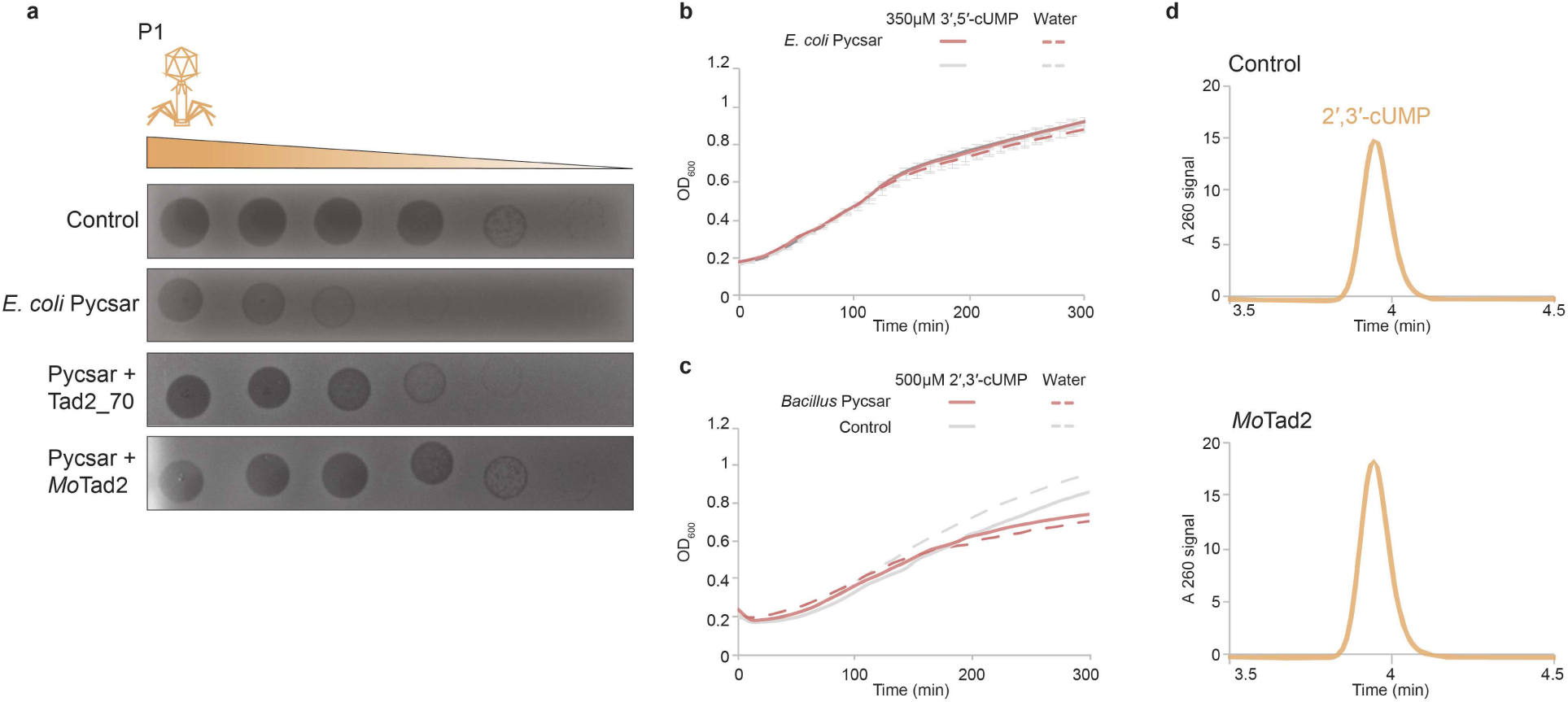
Specificity of Pycsar molecules. (A) Tad2 homologs (*Mo*Tad2 and Tad2_70) capable of overcoming Pycsar (cCMP) defense. Shown are tenfold serial dilution plaque assays, comparing the plating efficiency of phage P1 on bacteria that express Pycsar alone or with Tad2 homologs. Images are representative of three replicates. (B) cUMP does not inhibit the *E. coli* Pycsar. *E. coli* cells expressing the Pycsar system from *E. coli* E831 (naturally utilizing the signaling molecule cCMP), or control cells not expressing Pycsar, were grown in liquid media either with or without the addition of 350 µM cUMP. (C) cCMP does not inhibit the *B. subtilis* Pycsar. *B. subtilis* cells expressing the Pycsar system from *Bacillus* sp. G1 (naturally utilizing the signaling molecule 3′,5′-cUMP), or control cells not expressing Pycsar, were grown in liquid media either with or without the addition of 500 µM 2′,3′-cUMP. (D) HPLC analysis of 2′,3′-cUMP incubated with either purified GFP (control, top lane) or with purified *Mo*Tad2 at 1:4 molar ratio. The data indicate that *Mo*Tad2 does not sequester 2′,3′-cUMP.

**Figure S3.**
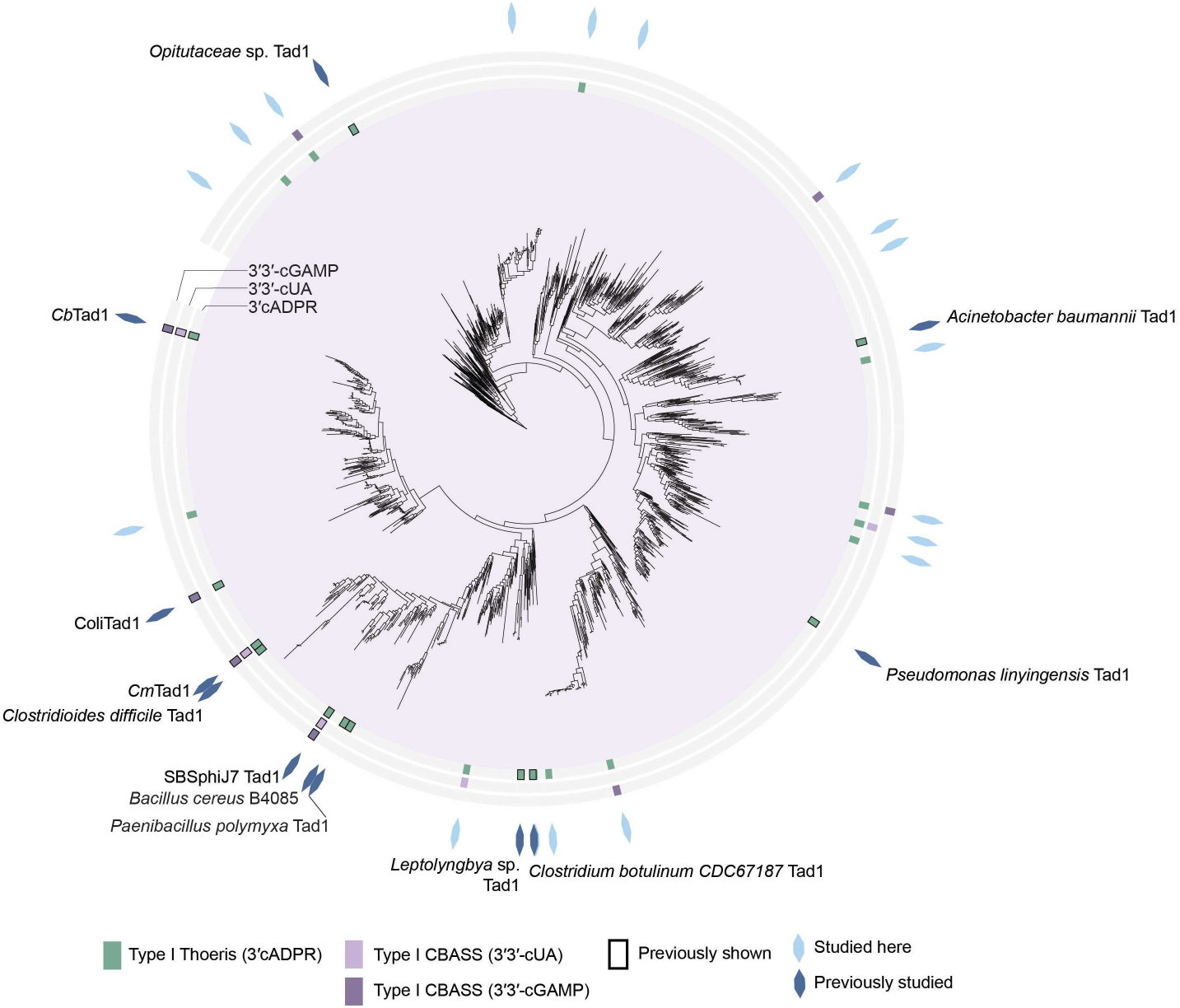
Phylogenetic analysis of Tad1 homologs. Shown is a phylogenetic tree of Tad1 homologs from phage genomic and metagenomic protein databases (n = 1,587 non redundant sequences). Previously studied homologs are marked by dark blue shapes on the outer rim of the tree; homologs tested in the current study are in light blue. Homologs that inhibit the defense system are marked with the respective color in the inner rings.

**Figure S4.**
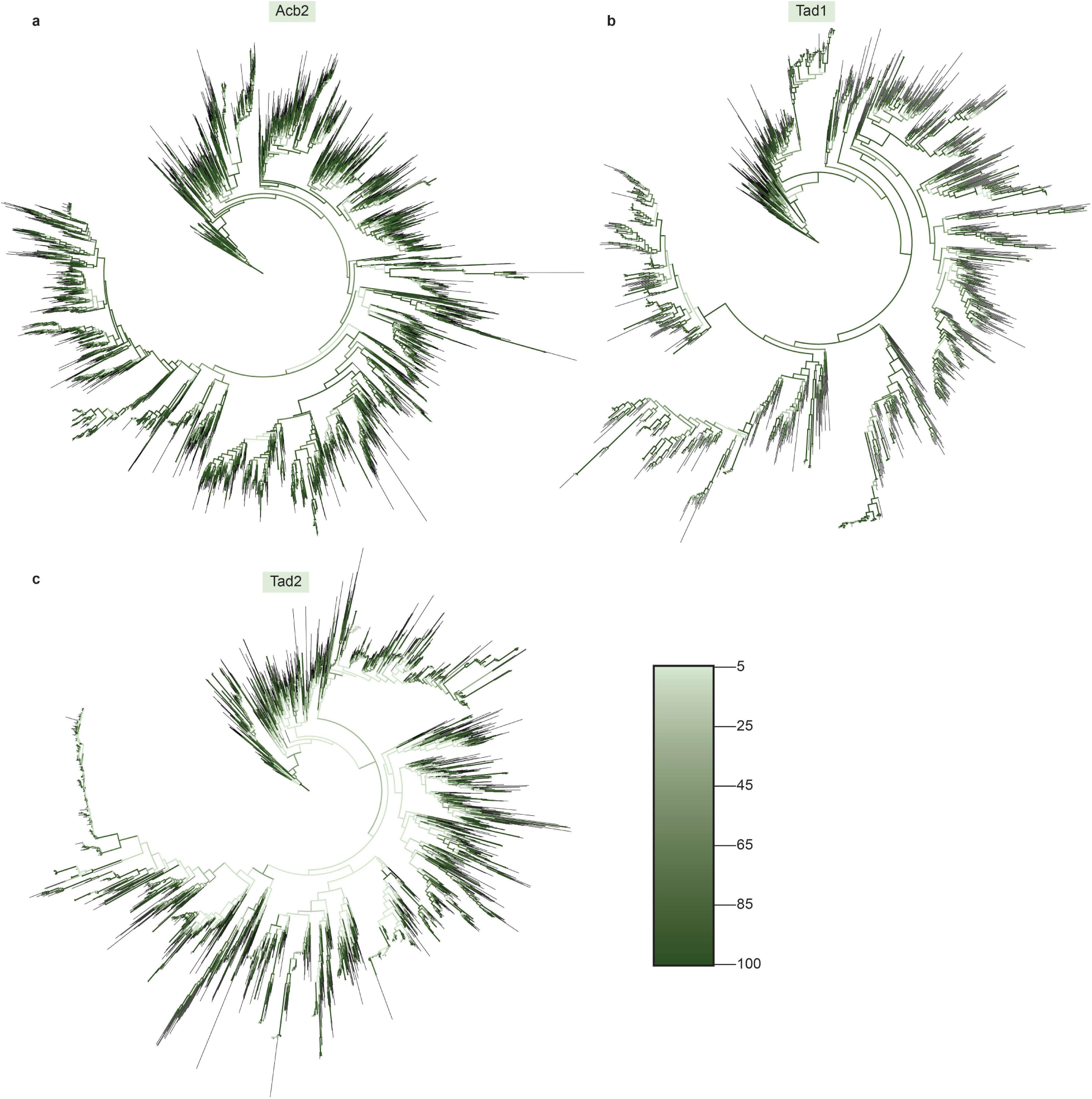
Phylogenetic trees colored by bootstrap values. Shown are the phylogenetic trees of Acb2 (A), Tad1 (B) and Tad2 (C). Tree branches are colored according to bootstrap values.

