## Supplementary material for "Functional diversity of phage sponge proteins that sequester host immune signals": Table S3

**Table S3. Crystallographic Statistics, Related to Figures 2**

|  | **Acb2 homolog 43–3′ADPR** |
| --- | --- |
| Resolution (Å)^a^ | 68.83–1.56 (1.59–1.56) |
| Wavelength (Å) | 0.97920 |
| Space group | P 1 2_1_ 1 |
| Unit cell: a, b, c (Å) | 60.12, 53.67, 73.48 |
| Unit cell: α, β, γ (°) | 90.00, 110.49, 90.00 |
| Molecules per ASU | 6 |
| Total reflections | 422,579 (19,721) |
| Unique reflections | 62,049 (3,073) |
| Completeness (%)^a^ | 99.1 (98.6) |
| Multiplicity^a^ | 6.8 (6.4) |
| *I/σI^a^* | 15.4 (1.7) |
| CC(1/2)^b^ (%)^a^ | 99.9 (82.1) |
| Rpim^c^ (%)^a^ | 2.8 (39.5) |
| Resolution (Å) | 68.83–1.56 |
| Free reflections | 2,942 |
| R-factor / R-free | 17.05 / 19.54 |
| Bond distance (RMS Å) | 0.009 |
| Bond angles (RMS °) | 1.090 |
| No. atoms: protein | 3,312 |
| No. atoms: ligand / ion | 70 |
| No. atoms: water | 556 |
| Average B-factor: protein | 22.65 |
| Average B-factor: water | 35.20 |
| Ramachandran plot: favored | 99.75% |
| Ramachandran plot: allowed | 0.25% |
| Ramachandran plot: outliers | 0.00% |
| Rotamer outliers | 0.28% |
| MolProbity^d^ score | 1.28 |
| Protein Data Bank ID | 9PTQ |

^a^ Highest resolution shell values in parentheses

^b^ (Karplus and Diederichs, 2012)

^c^ (Weiss, 2001)

^d^ (Chen et al., 2010)
